# Decoding medial entorhinal cortical dynamics produces planning-like alternations in hippocampal theta sequences

**DOI:** 10.64898/2025.12.29.696939

**Authors:** Masahiro Nakano, Caswell Barry, Claudia Clopath

**Author notes:** Correspondence to Claudia Clopath. These authors jointly supervised this work and contributed equally (Caswell Barry, Claudia Clopath).

## Abstract

The hippocampus is central to memory and spatial planning. A prominent candidate mechanism supporting navigational decision making is hippocampal theta sequences—brief (∼120ms) place-cell sequences representing future trajectories—which have been interpreted as planning signals based on their alternation between maze arms in T-maze tasks. However, recent work suggests that intrinsic left-right alternation in medial entorhinal cortex (MEC) theta sequences may underlie this phenomenon. Here, we test this hypothesis using a model of MEC theta dynamics in a T-maze and show that standard Bayesian decoding yields hippocampal theta sequences that alternate between arms. Analysis of simultaneous MEC-hippocampal recordings reveals tight coupling of MEC and hippocampus and coordinated left-right alternation. Notably, this alternation occurs not only at choice points but also during inbound and forced-turn trials. These findings suggest that left-right alternation in hippocampal theta sequences is a ubiquitous phenomenon driven by MEC dynamics, challenging the interpretation that these specifically reflect intentional decision-making or planning processes.

## Introduction

The hippocampus is essential for both remembering past experiences and using that information to guide future behavior. Damage to this structure causes profound memory deficits in humans and animal models (*1–3*), but its role extends beyond memory storage and retrieval. The hippocampus also enables imagination of future scenarios (*4*), something that is impaired in patients with hippocampal amnesia (*5*). This capacity for prospective thinking underlies planning, and planning-related representations have been identified in the human hippocampus (*6, 7*)

In rodents, the hippocampus exhibits a prominent 7–12 Hz theta rhythm that emerges from the combined activity of local interneurons and medial septal inputs (*8–10*). Individual hippocampal neurons fire preferentially near the trough of theta cycles (*11*). Place cells— neurons that fire when an animal occupies specific spatial locations—show systematic phase relationships with theta oscillations (*12–14*). When an animal traverses a place field, the corresponding place cell initially fires at late theta phases, then progressively earlier— a phenomenon known as phase precession (*12, 14*). At the population level, phase precession corresponds to rapidly progressing spatial trajectories within each theta cycle: nearby locations are represented early in the cycle while progressively distant locations are represented later. These “theta sequences” reflect temporally organized firing of place cell assemblies that sweep ahead of the animal’s current position (*11, 15, 16*). When Bayesian decoding is applied to these population patterns, the decoded position sweeps forward within each theta cycle, extending well beyond the animal’s actual location (*16–18*). It is widely held that theta sequences temporally compress spatial information into windows suitable for spike-timing-dependent-plasticity (STDP) (*14, 19, 20*), potentially supporting memory formation. Indeed, disrupting theta sequences impairs memory encoding (*18, 21*– *23*).

Theta sequences may also contribute to memory recall and planning (*24–27*). Since the initial report by Johnson and Redish (2007) (*28*), various studies have described theta-related representations that may reflect planning processes in rodents (*28–34*). In particular, Kay et al. (2020) (*30*) reported theta sweeps extending toward both the left and right arms as animals approached the decision point in a T-maze alternation task. Although these sweeps did not predict the eventual choice, they may represent the animal evaluating alternative options—a potential neural correlate of planning (*35, 36*). This finding aligns with the idea that theta oscillations may alternate between encoding and recall (*37*), representing the current and the future, or the real and the imagined (*38*).

The medial entorhinal cortex (MEC) is a major source of input to the hippocampus and is thought to play a key role in temporal coding within the hippocampal network (*39–44*). A variety of spatially tuned cell types have been identified in MEC, including grid cells, border cells, and object–vector cells (*45–48*). MEC activity is also strongly modulated by theta-band oscillations (*49, 50*), and grid cells exhibit phase precession (*13*). Recent work has demonstrated that MEC itself exhibits theta sequences (*50*). Notably, these theta sweeps progress forward but with a directional offset: MEC activity alternates between leftward and rightward sweeps across consecutive theta cycles. Moreover, hippocampal theta sweeps were found to alternate in a similar left–right pattern, temporally aligned with those in MEC. These observations raise the question of whether hippocampal left–right alternating theta sweeps observed in T-maze tasks reflect active planning within the hippocampus, or whether they simply mirror underlying MEC dynamics (as discussed in Vollan et al. (2025) (*50*)).

We tested this idea by modeling MEC and hippocampal theta sweeps in a T-shaped maze. Specifically, we simulated straight left–right alternating MEC sweeps that disregarded the physical T-shape and sampled spikes from place cells with fields on or near the T-maze. In this way, by applying Bayesian decoding to the population activity, we obtained hippocampal theta sweeps that appeared to traverse the left and right arms. Consistent with the left–right alternation in MEC, hippocampal sweeps also alternated between the two arms. We then turned to data analysis. Using a previously published dataset from Vollan et al. (2025) (*50, 51*), we found that hippocampal sweeps at the decision point alternated in coordination with MEC. Notably, this left–right alternation also occurred during inbound trials, where the correct choice was fixed, and even at the corner where the animal could only proceed in one direction. Hippocampal left-right alternation was always preceded by MEC representation. These findings support the view that left–right alternation of hippocampal theta sweeps is a ubiquitous phenomenon. We finally discuss experimental approaches to test more directly whether hippocampal theta sweeps constitute neural correlates of planning.

## Results

### Modeling left–right MEC sweeps reveals planning-like alternation in hippocampal activity

We set out to investigate whether left–right sweeps originating from MEC are sufficient to generate hippocampal representations that resemble alternating sweeps along the T-maze arms (Fig. 1A, B). To do so, we built a computational model consisting of an agent running down a T-maze, with the “MEC latent” sweeping left and right (Fig. 1C, left). During the early phase of theta, the MEC latent remained local, matching the agent’s actual position - during the late phase, it swept ahead of the agent’s position by an offset of 20 degrees from the head direction. The offset direction alternated between left and right in consecutive theta cycles (see Methods). This MEC latent drove activity in hippocampal place cells (Fig. 1A). Each hippocampal place cell was assigned a place field center uniformly distributed along the linearized track on the T-maze (Supp. Fig. 1A; left), and its instantaneous firing rate was determined by the Euclidean distance between the place field center and the MEC latent. Importantly, physical maze boundaries were ignored, so firing rate depended solely on this distance (see example firing rate map in Fig. 1C, right - note that the firing rate maps were calculated relative to the agent’s position, not the MEC latent).

**Fig. 1.**
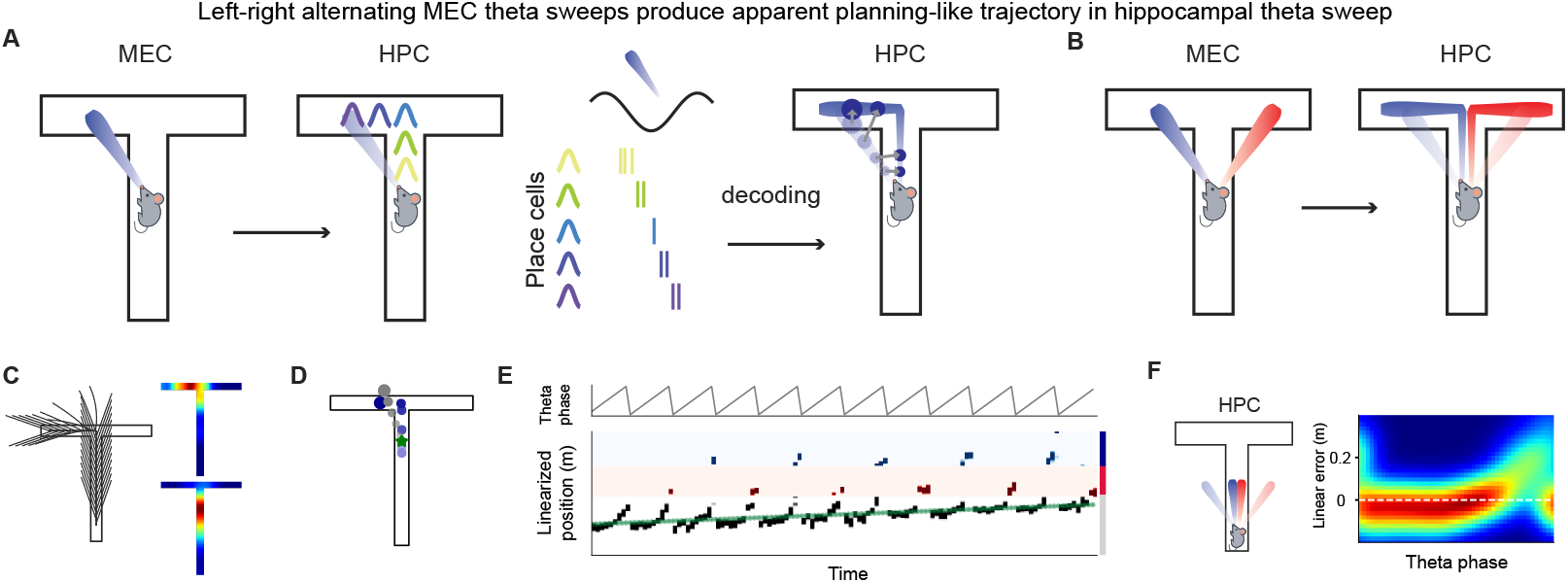
Modeling left–right MEC sweeps reveals planning-like alternation in hippocampal activity. (**A, B**) Model schematics showing hippocampal (HPC) sequences as projections of MEC sequences onto the T-maze. (**C**) Left: Alternating latent sequences used in simulation. Right: Two example hippocampal place cells. (**D**) Example theta sequence decoding showing current position (green), MEC latent (grey), and decoded position (blue), with opacity indicating temporal progression. (**E**) Decoded hippocampal sequences alternate between left (blue) and right (red) arms. Top: Theta phase. Bottom: Linearized likelihood map with darker colors showing higher likelihood; colors distinguish center, left, and right arms; green shows actual position. (**F**) Relationship between theta phase and linear error at the bottom of the center arm.

We next applied Bayesian decoding to infer latent position from place cell activity, using a maximum a posteriori (MAP) decoder. Because the firing rate map was only defined for locations the agent had visited, and decoding assigns the most likely position bin given the population activity, the decoder could not recover the true MEC latent, which sometimes extended beyond the T-maze (Fig. 1A). Instead, the decoder chose the best estimate within the area of the track. Consequently, decoding full MEC latent sweeps directed left while the agent was running in the center arm produced apparent sweeps that extended along the stem and into the left arm (Fig. 1D). Hippocampal theta sweeps decoded in the T-maze near the decision point showed significant alternation between left and right arms, reflecting the alternation in MEC latent (Shuffle test. See Methods for detail, Fig. 1E). We also examined theta sweeps when the agent was far from the decision point, effectively making the track linear. Consistent with the idea of forward left and forward right sweeps projecting onto the linear track, we observed forward-directed theta sequences (Fig. 1F). We then repeated the simulations under the assumption that some place cells had fields outside the physical T-maze. Place-field centers were uniformly distributed within the square environment that contained the T-maze. The square environment was designed so that all latent positions were contained within its boundaries. (Supp. Fig. 1A; right). In this case, too, left–right MEC sweeps produced hippocampal theta sweeps alternating between the two arms (Supp. Fig. 1A, B). Together, these results demonstrate that the combination of Euclidean left–right MEC sweeps and standard Bayesian decoding is sufficient to produce hippocampal theta sweeps that could be interpreted as neural substrates of planning, as originally suggested in Vollan et al. (2025)(*50*).

### Simultaneous MEC–hippocampal recordings reveal consistent coupling and left–right alternation at the T-maze decision point

If hippocampal left-right alternating sequences inherit their structure from MEC, then the timing and direction of sweeps in these regions should match. To test this possibility, we analyzed a publicly available dataset from Vollan et al. (2025)(*50*) containing simultaneous recordings from MEC and hippocampus while a rat performed an alternation task(*51*). They used an M-maze, a variant of the T-maze in which the left and right arms have an extra turn before the reward well. The dataset included 287 hippocampus cells and 681 MEC and para-subiculum (PaS) cells from a single animal. PaS exhibits a representation similar to that of MEC but shows stronger modulation by head direction (*50, 52, 53*). Vollan and colleagues reported that grid cells were predominantly found in MEC, whereas head-direction-modulated cells were more common in PaS, from the inferred anatomical location of the clusters (*50*). However, defining grid cells in a maze environment is challenging, partly due to limited spatial sampling and partly due to environmental influences on grid cell firing patterns (*54*). Since MEC and PaS directional signals are aligned (*50*), and our interest lies in the sweeping direction of MEC theta sequences, we included all units recorded in the MEC probe (MEC-Pas). Using Bayesian acausal population decoding, we first verified that hippocampus place cells exhibited forward sweeping theta sequences along the linear portion of the central arm (Supp. Fig. 2A). Decoding errors were lowest during the early phase of theta and increased during the late phase of theta, consistent with previous reports on theta sequence in linear tracks (*16–18*). We then examined activity as the animal approached the decision point (example outbound trial, Fig. 2A left). Clear alternation of theta sweeps toward the left and right arms was observed, replicating the original findings (Fig. 2A right, shuffle test, p<0.05; 1,000 permutations, all outbound trials combined (*30*)). Consistent with this population-level result, we identified a pair of place cells with place fields in the top-left and top-right corners of the maze (Fig. 2B left). This pair showed theta-cycling activity as the animal approached the decision point (Fig. 2B middle. Same trial as Fig. 2A). Cross-correlation analysis across all outbound trials confirmed consistent cycling between the two cells (Fig. 2B right). Next, we examined MEC-PaS representations when hippocampal sweeps alternated between the left and right arms. We found MEC cells that were active immediately before hippocampus place cells (Fig. 2C middle). These MEC cells were modulated by head direction, and their preferred direction matched the direction of the place fields of hippocampal cells that followed the activity (Fig. 2D).

**Fig. 2.**
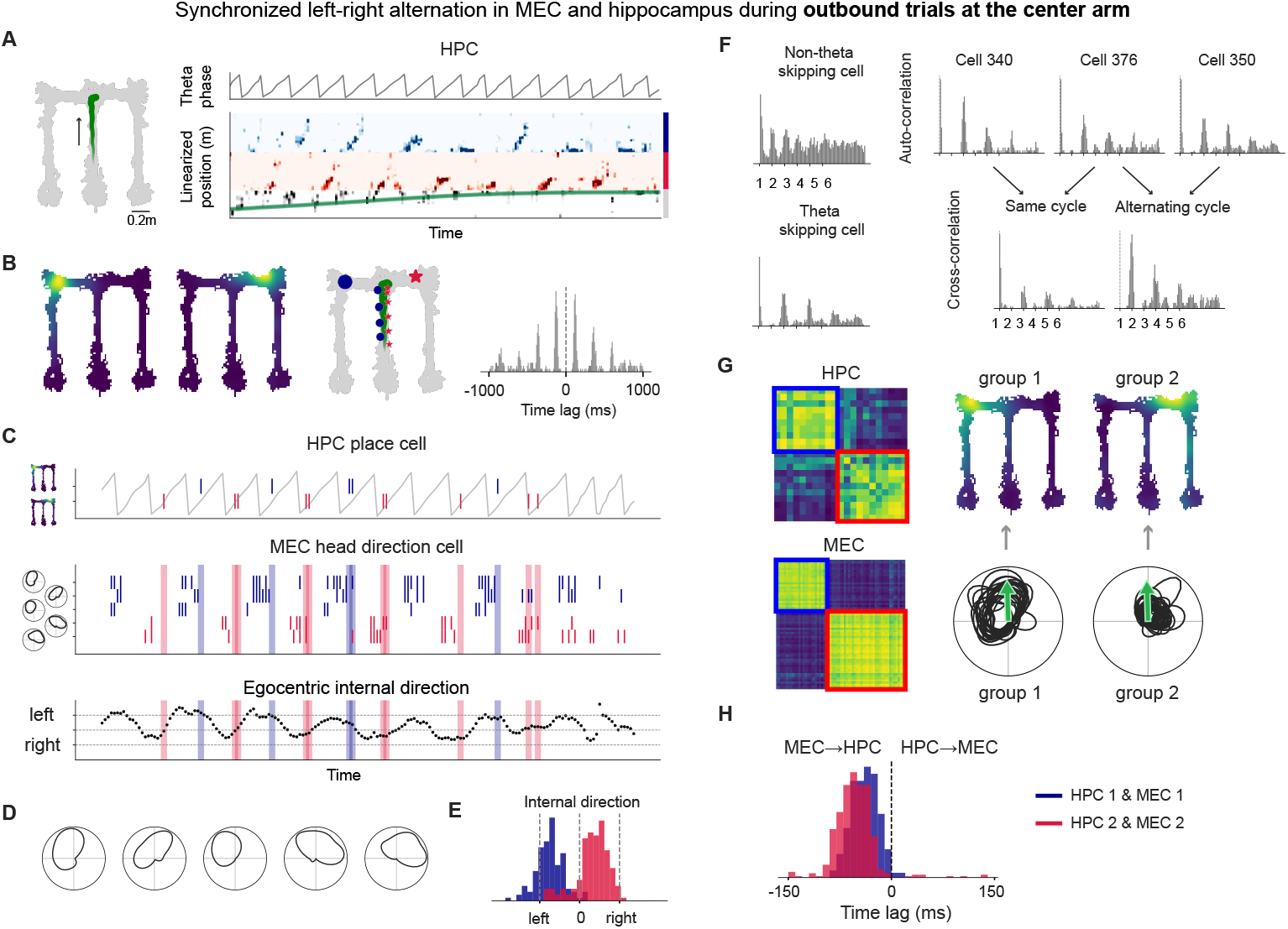
Simultaneous MEC–hippocampal recordings reveal consistent coupling and left–right alternation at the T-maze decision point. (**A**) Example outbound trial showing the animal’s trajectory (left), theta phase (top right), and population decoding likelihood (bottom right). (**B**) Left: Firing rate maps of two hippocampal cells. Middle: Example outbound trial showing the trajectory (green) and the animal’s positions when the two hippocampal cells were active (left; blue, right; red). Large blue circle and red star indicate the place-field centers. Right: Cross-correlogram between the two hippocampal cells across all outbound trials. (**C**) Same trial as in (B), showing activity of hippocampal cells (top), MEC cells (middle), and egocentric internal direction– defined as the difference between head direction and MEC internal direction– (bottom). (**D**) Head direction tuning curves of the MEC cells shown in (C). (**E**) Distribution of internal directions when the two hippocampal cells were active; colors correspond to panel (B). Internal direction of 5 frames before the place cell active is plotted, to reflect the time lag between MEC and hippocampus. **(F)** Example auto-correlograms of a non-theta skipping cell and a theta skipping cell (left), and cross-correlograms of cell pairs active in the same versus alternating theta cycles (right). **(G)** Theta-skipping cell analysis showing clustering (left), summed firing-rate maps for hippocampal cells (top middle), and head-direction tuning for MEC cells (bottom middle). **(H)** Distribution of cross-correlation lags for hippocampus-MEC cell pairs with matching direction preference.

Vollan and colleagues computed an “internal direction” signal from MEC and PaS activity and showed that, as in open-field recordings, this internal direction alternated between left and right on the M-maze (see Supp. Fig. 8 in (*50*)). We found that, similar to the open-field results, the left-right alternation of MEC egocentric internal direction, defined as the difference between head direction and MEC internal direction, was consistent with the relative direction of hippocampus place field with respect to head direction (Fig. 2C bottom, 2E). Previous reports suggest that hippocampal activity lags behind MEC activity, reflecting the information flow from MEC to hippocampus (*50, 55, 56*). Consistently, place cell activity occurred during the late phase of internal-direction switching (Fig. 2C bottom).

To systematically quantify this phenomenon, we analyzed spike-time correlations. Many hippocampal and MEC neurons are modulated by the theta rhythm (*12, 49, 57, 58*), resulting in autocorrelation peaks every 100–120 ms (non-theta-skipping cells; Fig. 2F top left). In contrast, some cells exhibit peaks separated by two theta cycles (theta-skipping cells; Fig. 2F bottom left). Theta-skipping cells have been reported in the hippocampus, MEC, thalamus, and PFC (*34, 49, 50, 59*) and may contribute to alternating theta sweeps. Notably, theta skipping can occur only in specific spatial contexts. For instance, theta skipping in a given place cell might only be obvious when the animal is running toward the place field center, but this effect is obscured when all time points are included (Supp. Fig. 2B). We therefore identified theta-skipping cells within spatially and directionally restricted time windows. First, we restricted the analysis to strongly theta-skipping cells. Across 43 outbound trials, we detected 21 theta-skipping hippocampal cells (Fig. 2F, top right; see Methods for details). We then computed cross-correlations for all identified theta-skipping cell pairs. Cell pairs with peaks at even theta cycles fired within the same theta cycle, whereas those with peaks at odd theta cycles fired in alternating cycles (Fig. 2F, bottom right). From these cross-correlations, we derived a “theta-skipping index” for all pairs and constructed a theta-skipping index matrix (See Methods). Clustering this matrix revealed two major hippocampal populations (Fig. 2G top left, Methods), corresponding to top-left and top-right place cells (Fig. 2G top right; Sum of firing rate map of all cells in each group. Individual maps in Supp. Fig. 2C). Firing rate maps had significantly higher correlation within the group compared to across groups (shuffle test, p<0.05; 5,000 permutations, Methods). Applying the same analysis to MEC theta-skipping cells again yielded two groups, corresponding to left- and right-direction-modulated cells (Fig. 2G bottom). Similarly, the overlap of the head direction tuning curve was higher within the group compared to across groups (shuffle test, p<0.05). Finally, we measured the peak time lag between MEC and hippocampal groups. Left- and right-direction-modulated MEC cells consistently preceded the corresponding top-left and top-right hippocampal place cells, with an average lag of ∼45ms (group 1; -43.0ms, t(249)=-35.6, p<10^-20^, group 2; -48.9ms, t(417)=-21.2, p<10^-20^, one-sample t-test, Fig. 2H). We repeated this analysis with looser criteria for theta-skipping cells. In this case, cells that demonstrated weaker but significant theta-skipping were included (see Methods and Supp. Fig. 2E). These cells often showed a local peak in one theta cycle apart, but the peak size was comparable to the second theta cycle peak, which is different from a classical decay over theta cycles in theta-modulated cells. These contained place cells that had fields in the center arm, but had stronger coupling to the either arm place cells. With this criterion too, theta skipping cells in hippocampus and MEC were both clustered into two groups, which corresponded to relative left and right direction, and MEC-hippocampal coupling was also preserved (shuffle test, hippocampal firing rate map correlation; p<0.05, MEC head direction tuning curve correlation; p<0.05, time lag between MEC and hippocampus; group 1;-30.9ms, t(944)=-22.0, p<10^-20^, group 2; -35.2ms, t(1215)=-26.2, p<10^-20^). In summary, within the same dataset, we show that (1) hippocampal theta sweeps alternate between left and right arms near the decision point, and (2) this alternation is preceded by a matching left-right representation in MEC.

### MEC-hippocampal coupling occurs under less cognitively demanding conditions

In alternation tasks, neural representations have predominantly been studied at the decision point, where cognitive demand is highest and planning is most relevant. However, MEC left–right alternations occur ubiquitously across the M-maze(*50*). Building on the idea that hippocampal theta sweeps may reflect MEC sweeps, we asked whether MEC–hippocampal synchronization and left–right alternation extend beyond the decision point. We focused on two additional conditions, beginning with the inbound trials. During inbound trials, when the animal returns from either arm, the task rule requires it to always turn into the center arm– thus demanding less planning or decision-making. Indeed, animals learn inbound trials more rapidly than outbound trials (*60*), and perturbations that impair learning have a smaller effect on inbound trials (*60, 61*). If hippocampal theta sweeps are closely tied to cognitive demands or planning states, they might be biased toward the correct choice, or show reduced sampling-like in these low-demand trials. We examined inbound trials in which the animal returned from the left arm. Similar to the outbound center-arm trials, we observed alternating theta sequences that swept toward the center and right arms (Fig. 3A). Consistent with this, a hippocampal place-cell pair showed alternating activity (Fig. 3B), with place fields centered in the right and center arms, respectively. In the MEC, we identified direction-selective cells tuned to rightward and downward movements that were active immediately before the corresponding hippocampal place cells (Fig. 3C, D). Egocentric internal direction was also consistent with the direction of the hippocampal place fields (Fig. 3E). Applying the same theta-skipping analysis revealed two distinct cell groups in both hippocampus and MEC (Fig. 3F left). Each hippocampal and MEC group exhibited correlated place fields and head-direction tuning curves, respectively (Fig. 3F right, shuffle test; hippocampus, p<0.05, MEC, p<0.05). Moreover, MEC groups with egocentric direction selectivity matching the downstream hippocampal place fields consistently preceded hippocampal activity (Fig. 3G, group 1; t(43)=-27.5, p<10^-20^, group 2; t(179)=- 17.3, p<10^-20^). Similar results were obtained using looser criteria for theta skipping cells (shuffle test; hippocampus, p<0.05, MEC, p<0.05, time lag; group 1; t(293)=-19.1, p<10^-20^, group 2; t(454)=-13.3, p<10^-20^). Similar results were obtained for inbound trials from the right arm (Supp. Fig. 3B, strict threshold; shuffle test; hippocampus, p<0.05, MEC, p<0.05, time lag; group 1; t(109)=-11.5, p<10^-20^, group 2; t(68)=-7.4, p=1.4×10^-10^).

**Fig. 3.**
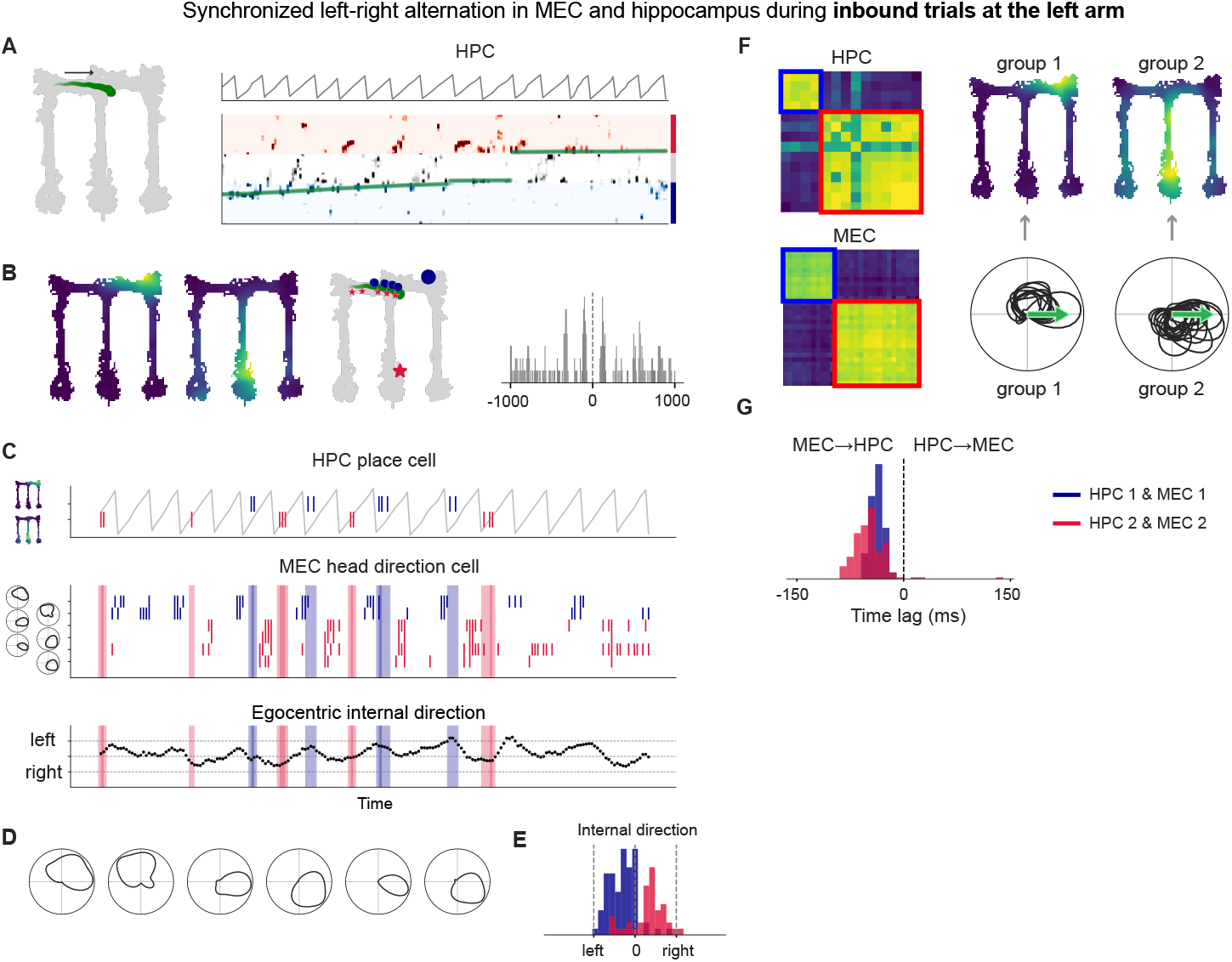
MEC-hippocampal coupling occurs under less cognitively demanding conditions. (**A**) Example inbound trial when the animal was running toward the junction from the left arm, showing the animal’s trajectory (left), theta phase (top right), and population decoding likelihood (bottom right). (**B**) Left: Firing rate maps of two hippocampal cells. Middle: Example inbound trial from the left arm showing the trajectory (green) and the animal’s positions when the two hippocampal cells were active (top-right; blue, center arm; red). Large blue circle and red star indicate the place-field centers. Right: Cross-correlogram between the two hippocampal cells across all inbound trials from the left arm. (**C**) Same trial as in (B), showing activity of hippocampal cells (top), MEC cells (middle), and egocentric internal direction– defined as the difference between head direction and MEC internal direction– (bottom). (**D**) Head direction tuning curves of the MEC cells shown in (C). (**E**) Distribution of internal directions when the two hippocampal cells were active; colors correspond to panel (B). Internal direction of 5 frames before the place cell active is plotted, to reflect the time lag between MEC and hippocampus. (**F**) Theta-skipping cell analysis showing clustering (left), summed firing-rate maps for hippocampal cells (top middle), and head-direction tuning for MEC cells (bottom middle). (**G**) Distribution of cross-correlation lags for hippocampus-MEC cell pairs with matching direction preference.

### MEC-hippocampal coupling persists even at forced turn corners

To study how ubiquitous MEC and hippocampus coupling is, we took advantage of the M-shaped design of the maze, which introduced a second turning corner absent in a T-maze. This corner is typically not considered a decision point, as there is only one possible path. Consequently, cognitive demand should be lowest here compared to either the outbound or inbound decision points. We analyzed neural representations when the animal ran from the center arm toward the top-left corner (Fig. 4A). In most trials (15/20), the animal made a smooth turn at the corner and continued down the left arm, while in a subset of trials (4/20), it briefly visited the top-left corner before descending (Supp. Fig. 4A). Surprisingly, we identified a hippocampal place-cell pair exhibiting theta cycling (Fig. 4B). The place fields of these two cells corresponded to the left arm and the top-left corner, which aligned with the animal’s leftward and rightward directions, respectively. We found MEC cells that were selective to bottom left and top left direction, that were active immediately before left arm and top-left corner place cells, respectively (Fig. 4C, D). Egocentric internal direction was consistent with the timing of place cell activity (Fig. 4E). Applying the same analysis as before, we again identified two groups in both hippocampus and MEC: one corresponding to the top-left corner and one to the left arm (Fig. 4F). The MEC groups showed leftward and rightward direction selectivity and consistently preceded the activity of the matching hippocampal groups (group 1; t(71)=-12.7, p<10^-20^, group 2; t(44)=-5.02, p=4.6x10^-6^, Fig. 4G). Kay et al. (2020) reported theta-skipping cell pairs in a similar condition, noting that these pairs consisted of cells with outbound- and inbound-dominant responses. Our results, derived from large-scale recordings, offer an alternative interpretation that emphasizes the spatial location of place fields rather than directional dominance.

**Fig. 4.**
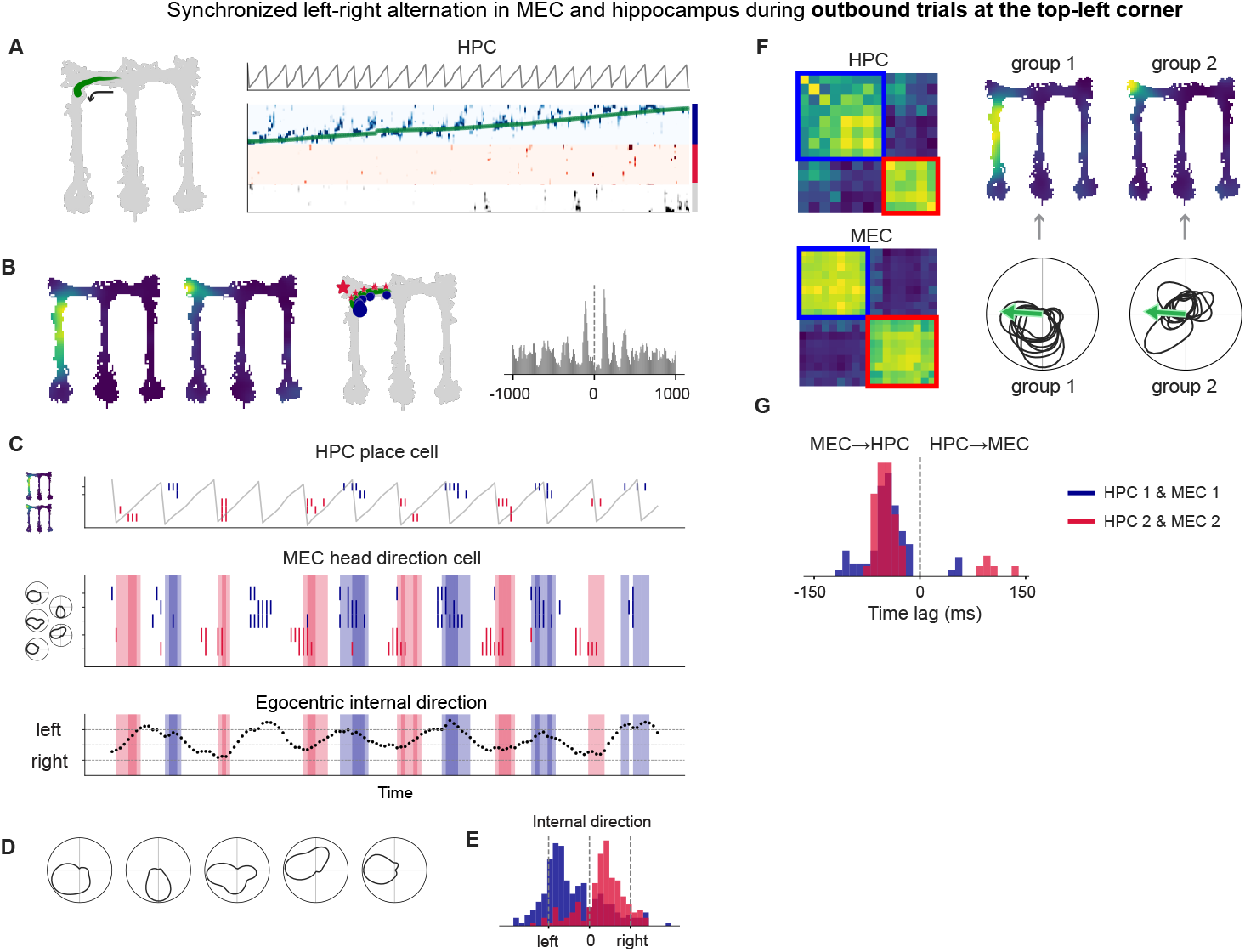
MEC-hippocampal coupling persists even at forced turn corners. (**A**) Example outbound trial when the animal was running toward the top-left corner from the junction, showing the animal’s trajectory (left), theta phase (top right), and population decoding likelihood (bottom right). (**B**) Left: Firing rate maps of two hippocampal cells. Middle: Example outbound trial towards the left arm showing the trajectory (green) and the animal’s positions when the two hippocampal cells were active (inner corner; blue, top-left corner; red). Large blue circle and red star indicate the place-field centers. Right: Cross-correlogram between the two hippocampal cells across all outbound trials towards the left arm. (**C**) Same trial as in (B), showing activity of hippocampal cells (top), MEC cells (middle), and egocentric internal direction– defined as the difference between head direction and MEC internal direction– (bottom). (**D**) Head direction tuning curves of the MEC cells shown in (C). (**E**) Distribution of internal directions when the two hippocampal cells were active; colors correspond to panel (B). Internal direction of 5 frames before the place cell active is plotted, to reflect the time lag between MEC and hippocampus. (**F**) Theta-skipping cell analysis showing clustering (left), summed firing-rate maps for hippocampal cells (top middle), and head-direction tuning for MEC cells (bottom middle). (**G**) Distribution of cross-correlation lags for hippocampus-MEC cell pairs with matching direction preference.

This pattern was replicated across most corners of the maze, including the top-left inbound, top-right outbound, and top-right inbound directions (Supp. Fig. 4B–D). In the top-right corner inbound condition, there were fewer theta-skipping cells that met the criteria, and the overlap of firing-rate maps of hippocampal cells was non-significant (shuffle test, p=0.111). However, visual inspection suggested the tendency for hippocampal cell groups following left- and right-direction MEC cell activity to have relatively left- and right-directed place fields, respectively (Supp. Fig. 4D right). Thus, MEC–hippocampal synchronization and left–right alternation occur ubiquitously throughout the maze environment, extending beyond classical decision points.

## Discussion

Using a computational model, we demonstrated that combining left–right alternating theta sweeps with standard Bayesian decoding can produce an apparently planning-like representation in the hippocampus. Analysis of an existing dataset further revealed that during left–right alternating hippocampal theta sweeps in the T-maze, medial entorhinal cortex (MEC) representations are directionally aligned with, and critically, precede hippocampal activity—indicating robust MEC–hippocampal coupling. Importantly, this synchronization persists beyond the classic decision point, extending to cognitively less demanding inbound trials—where the correct turn is always predetermined—and even to maze corners, where no choice is required. These findings extend previous reports of MEC– hippocampal coordination in open-field environments by demonstrating that such synchronization also occurs during structured maze tasks and active behavior. Finally, we observed that even when the environment is designed to be “one-dimensional,” the rat brain represents it in a two-dimensional manner (Fig. 4). This suggests that the common practice of linearizing trajectories in hippocampal studies may obscure important aspects of spatial coding and directional representation.

Recent experiments have adopted environments with greater spatial freedom to study theta sequence representations, such as complex mazes (*31, 33, 62*) or open-field arenas (*32*). Comrie et al. (2024)(*33*) reported theta representations that were flexibly modulated by task rules and rewards that changed within a session, including representations of physically remote locations–features that are difficult to reconcile with the Euclidean framework of MEC representations thought to be inherited by the hippocampus. Vollan et al. (2026)(*63*) reported theta sweeps in MEC and PaS that were modulated by behaviorally salient directions. However, it is important to note that the direction of these theta sweeps was stable over several theta cycles, indicating that this phenomenon is distinct from the theta sweeps observed in T-maze tasks, which alternate on every theta cycle in both hippocampus and MEC. Yu et al. (2025)(*31*) and Tang et al. (2025)(*32*) similarly reported theta sweep directions biased toward goals, and both studies used attractor network models incorporating an egocentric goal signal. Importantly, both models relied on short-term firing rate adaptation to generate theta sweeps (*64, 65*), consistent with treating the MEC as a fully two-dimensional Euclidean system. However, it remains unclear how the same system could produce meaningful theta sweeps in the M-maze, where, during outbound trials in the center arm, the goal location is behind the animal. Overall, theta-related representations ranging from simple left–right alternation to more complex, task-modulated patterns have been reported. This suggests that theta activity may encompass multiple forms of representations, some of which could support planning. Most importantly, future experiments addressing this question must consider that seemingly non-cognitive dynamics in MEC could give rise to planning-like signals, potentially confounding interpretations, and should be designed accordingly.

To disentangle these possibilities, we propose an “F-maze” (Fig. 5). Topologically, it is identical to the T- or M-maze but differs in its physical layout: instead of two choice arms positioned left and right of the center arm, the F-maze places both arms on one side (in this case, the right). This design would allow us to test two key questions:

**Fig. 5.**
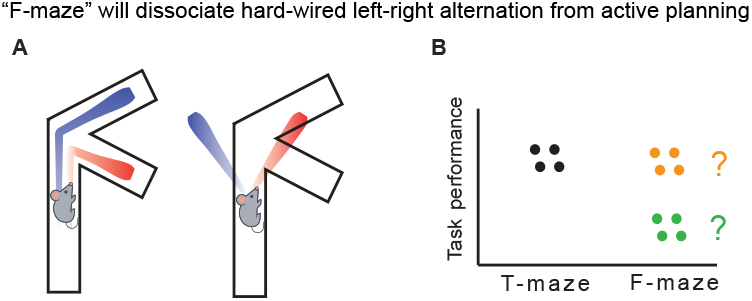
“F-maze” will dissociate hard-wired left-right alternation from active planning. (**A**) Schematics showing two possibilities of theta sweeps in the F-maze. Theta sweeps may alternate between two choice arms (left) or alternate between left and right (right). (**B**) Schematics showing two possibilities of task performance in the F-maze. Animals may show lower performance, potentially linked to theta sweeps (light green), or show a similar level of performance (orange).

1. Do we still observe alternating theta representations across the two choice arms?
2. Does behavioral performance remain comparable to that in the T-maze, or does it become more difficult for the animals?

If alternating theta representations are observed between the two right-side arms, this would argue against the idea that hippocampal left–right theta sweeps merely reflect intrinsic MEC dynamics (Fig. 5A). Regardless of whether MEC, hippocampus, or an upstream region drives the effect, such results would provide strong evidence that theta sweeps are modulated by task content. Conversely, if alternating representations are absent in the F-maze, two interpretations are possible. One is that left–right sweeps are a byproduct of MEC dynamics, but the brain can nevertheless use this representation for planning in downstream regions. In this case, performance in the T-maze may be higher because these representations aid planning, whereas performance in the F-maze would decline due to the loss of supporting information, or at least be different since animals need to use alternative mechanisms. Alternatively, if hippocampal theta sweeps are not used for planning—either within hippocampus or downstream areas—performance should remain comparable between maze types. In this scenario, the brain might instead rely on “splitter cells,” found in multiple regions including hippocampus, prefrontal cortex, entorhinal cortex, and thalamus (*66–68*) .

Finally, if theta is not primarily or solely used for memory recall or planning, what is its functional role? Theta’s involvement in encoding is increasingly well established, as perturbation studies have demonstrated memory impairments following theta disruption (*69–71*). Quirk et al. (2021) (*69*) perturbing the medial septum impaired performance only when applied during the return arm, not during the outbound center arm. Similarly, Zhang et al. (2021) (*70*) found that medial septum stimulation during the outbound center arm had no effect on performance, whereas stimulation in the delay area reduced ripple rates and task accuracy. These findings suggest that theta oscillations are involved in memory encoding, whereas retrieval may rely on replay mechanisms. Future experiments should therefore employ carefully designed behavioral tasks combined with causal manipulations to determine whether or to which extent theta also contributes to memory recall and planning.

In summary, our computational model confirmed the hypothesis that left–right sweeps in the medial entorhinal cortex (MEC) can give rise to planning-like activity in the hippocampus. Analysis of simultaneous MEC–hippocampal recordings revealed that their coupling occurs ubiquitously during maze alternation behavior, supporting the idea that hippocampal left–right sweeps may arise from MEC-driven dynamics.

## Methods

### Simulation of left-right sweeps

To simulate left-right alternating theta sweeps and sample place cell spikes, we used RatInABox (*72*). RatInABox accepts *Agents* that move around in an *Environment*. We created a T-maze *Environment* where the *Agent* had a predefined fixed behavior, starting from the bottom of the center arm, then ran towards the left arm, and the right arm alternately. We then modeled MEC as a *SubAgent* that inherited the behavioral parameters of the simulated *Agent*. The instantaneous theta phase was computed as a function of time and the simulated theta frequency. During the initial fraction of each theta cycle (parameter *α*), the *SubAgent’s* position coincided with that of the *Agent*. During the remaining fraction (1 − *α*), the *SubAgent* advanced away from the *Agent’s* position at a constant speed of *v*_*sub*_. The movement direction was offset by 20° relative to the *Agent’s* instantaneous head direction, consistent with experimental observations reported in (*50*). Place-cell populations were defined under two configurations. In the first configuration, place cells were distributed uniformly along the linearized T-maze trajectory at a constant spatial density (see Table 1). In the second configuration, a two-dimensional square environment was defined by embedding the T-maze within a surrounding padding region. Place-cell centers were positioned on a regular mesh grid at the specified areal density (Table 1). The place cell’s instantaneous firing rate was calculated based on the distance between the assigned place field center and the latent position (See Supp. Fig. 1 for the distribution of place field centers). The firing rate *r* was calculated as

**Table 1.** Parameters used in the simulation.

|  | Parameter name |  | Value | Unit |
| --- | --- | --- | --- | --- |
| Behavior | Running speed | | 0.3 | $\text{ms}^{-1}$ |
| Maze | T-maze size | Center arm length | 1 | m |
|  |  | Left and right arm length | 0.5 | m |
|  |  | Arm width | 0.1 | m |
| General | Time step size ($\Delta t$) | | 1 | ms |
|  | Simulation length |  | 20 | s |
| MEC theta sweeps | Theta sequence forward sweep fraction ($\alpha$) | | 0.5 | |
| | Theta sequence speed ($v_{sub}$) | | 8 | $\text{ms}^{-1}$ |
|  | Theta frequency |  | 10 | Hz |
| Place cells | Density of place cells for simulation 1 | | 500 | $\text{m}^{-1}$ |
| | Density of place cells for simulation 2 | | 2500 | $\text{m}^{-2}$ |
| | Place field width ($\sigma$) | | 0.15 | m |
| | Peak firing rate ($r_{max}$) | | 20 | Hz |

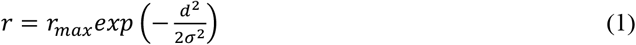

where *r*_*max*_ is the peak firing rate, *d* is the distance between the place field centre and the MEC latent, and *σ* is the place field width.

Then, binary spikes were sampled with a probability of *ρ*

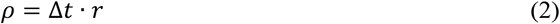

Where Δ*t* is the time step size.

See Table 1 for parameters used in the simulation.

### Path linearization

To linearize positions, six anchor points were defined corresponding to the center bottom, center top (junction), top left, bottom left, top right, and bottom right of the maze. Path points were then interpolated between these anchors, and all behavioral positions were projected onto the nearest path points.

### Temporal binning

In the dataset from Vollan et al. (2025), behavioral measures and neural activity were sampled in 10-ms bins. To match this, we resampled the simulated dataset at 10-ms bins.

All analyses were performed on this 10-ms binned data.

### Firing rate map / Head-direction tuning curve

Place fields were estimated in two coordinate spaces. On the linearized path, firing rates were binned at 2 cm resolution and smoothed with a 5 cm Gaussian kernel. Smoothing was constrained within each arm and did not cross boundaries. Linearized fields were used for population decoding in both simulations and data analyses. In two dimensions, the maze was divided into 2 cm × 2 cm bins, and kernel density estimation was applied with a 5 cm bandwidth. These two-dimensional fields were used for visualization and place-field correlation analyses (Figs. 2–4).

Circular kernel density estimation was performed using 6° bins and smoothed with a von Mises kernel (*κ* = 10).

In all cases, spike density and position density were calculated separately, and tuning curves were calculated by dividing spike density by position density.

### Population decoding

Population activity was decoded using a Bayesian approach with a uniform prior. Only frames with more than one spike were used in the analysis.

To quantify left–right alternation in the decoded trajectory, we identified time points when the decoded position lay in the left or right arm, respectively. These time series were treated analogously to spike trains. Following the procedure for theta skipping, we computed a skipping index for the left and right latent event trains (see *Theta-skipping index*). Theta-cycle indices were shuffled 1,000 times to generate a null distribution. The 5th percentile of this distribution defined the chance level of theta skipping; observed values below this threshold were considered to exhibit left–right alternation.

### Theta-skipping index

For correlation analyses, spikes within each 10ms bin were binarized. For each spike auto- or cross-correlogram, two peak values were extracted to quantify firing in alternating theta cycles:

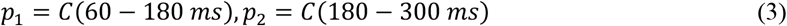

where *C*(*t*_1_ − *t*_2_) denotes the mean spike count within the specified time window [*t*_1_, *t*_2_]. *p*_1_ is the mean count of the auto- or cross-correlogram during the time window that corresponds to the next theta cycle, and *p*_2_ is for the second next theta cycle.

The theta-skipping index (TSI) was computed as

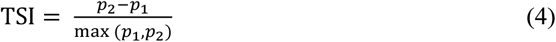

A value of 1 indicates complete theta skipping, whereas -1 indicates firing one theta cycle apart, reflecting theta cycling.

### Theta-skipping cells

To generate shuffled controls, spike times were randomized while preserving theta-phase structure. Theta cycles were first segmented, and their indices were then shuffled so that theta modulation was maintained after shuffling.

Cells were classified as “loose” theta-skipping if they met two criteria. First, mean firing rates exceeded a minimum threshold (hippocampus: 1 Hz; MEC: 5 Hz). Second, the observed TSI exceeded the 95th percentile of a null distribution generated from 1,000 shuffles.

For a stricter classification, cells were additionally required to have *TSI* > 0.5, excluding weak alternators (Supp. Fig. 2E).

### Clustering of theta-skipping cells

An affinity matrix was constructed using pairwise theta-skipping indices. Spectral clustering was applied with cluster numbers ranging from 2 to 8, and silhouette scores were computed. The cluster number yielding the highest score was selected.

### Firing-rate map and head-direction tuning correlation

To assess whether hippocampal firing-rate maps and MEC head-direction tuning curves were more similar within clusters than across clusters, we computed pairwise correlation matrices for all cells. The difference between the mean within-cluster and across-cluster correlation coefficients defined an index of within-cluster similarity. This index was recalculated 5,000 times with shuffled cluster labels to obtain a null distribution, from which p-values were derived.

### Time-lag analysis

For each hippocampal–MEC cell pair, cross-correlations were computed within a ±150 ms window. The lag corresponding to the peak correlation was taken as the pair’s time lag. After pooling across pairs, a one-sample t-test was performed against a mean of 0 to test for systematic lead–lag relationships

## Statistical Analysis

Data analyses were performed with custom-written scripts in Python, and clustering analyses and statistical analyses were conducted using the python package scipy and scikit-learn.

## Acknowledgments

We thank A.Z. Vollan, R.J. Gardner, M.-B. Moser, E.I. Moser for making the dataset and analysis code available and easy to use. We thank L. Frank for helpful comments.

## Author contributions

Conceptualization: MN, CB, CC

Methodology: MN

Investigation: MN

Visualization: MN

Supervision: CB, CC

Writing—original draft: MN

Writing—review & editing: MN, CB, CC

## Competing interests

Authors declare that they have no competing interests.

## Data and materials availability

Data analyzed in this paper is available at EBRAINS, https://doi.org/10.25493/R5FR-EDG. Code for reproducing the simulation and analyses in this article will be available at Zenodo https://doi.org/10.5281/zenodo.18340856

---

**Fig. S1.**
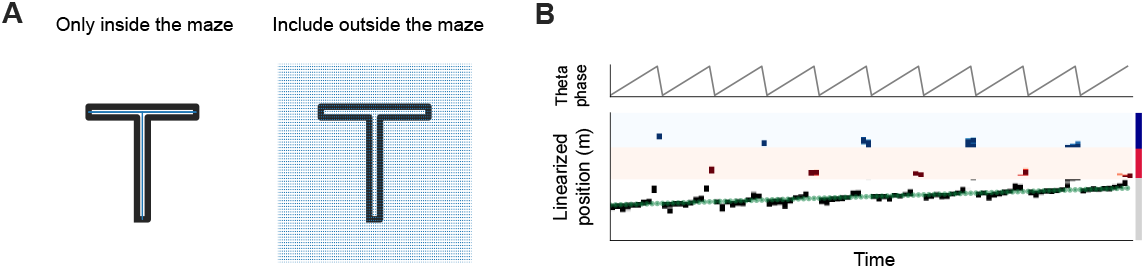
**(A)** Distribution of simulated place cells under two assumptions: one in which place cells are confined to locations within the maze, and another in which they also exist outside the maze. (**B**) Population decoding result for the condition where place cells extended beyond the maze boundaries.

**Fig. S2.**
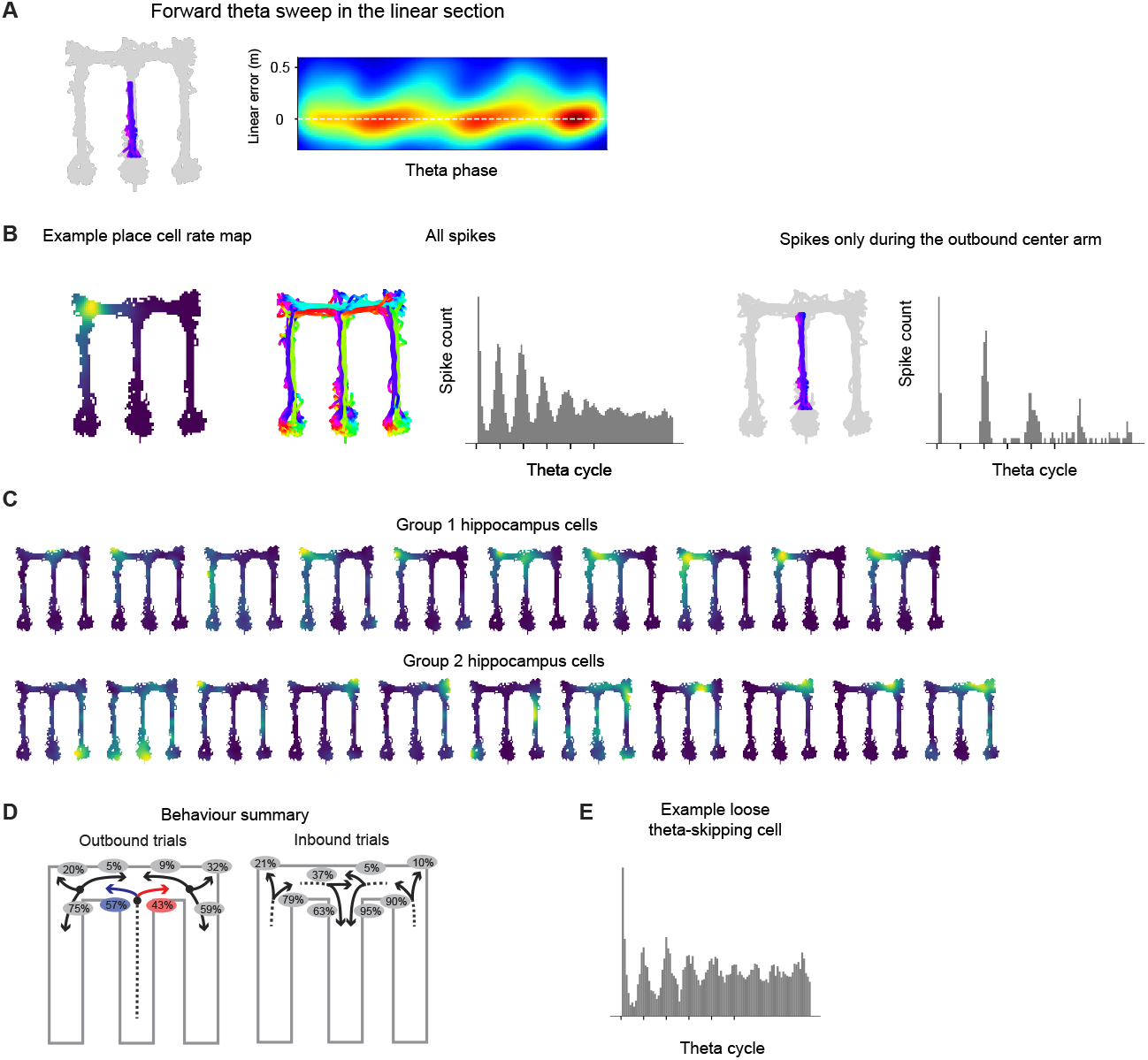
(**A**) Forward theta sweep in the center arm. Left: Animal’s trajectory corresponding to the analyzed timepoints. Right: Decoded population activity during these time points. (**B**) An example hippocampal cell showing strong theta skipping restricted to a specific region. Left: Firing-rate map of the cell. Middle: Using all time points. Right: Using time points when the animal was running outbound in the center arm. (**C**) Individual firing-rate maps of hippocampus cells belonging to the identified cluster. (**D**) Summary of the animal’s behavior during the analyzed sessions. (**E**) An example cell that satisfied the loose but not the strict criteria for theta skipping.

**Fig. S3.**
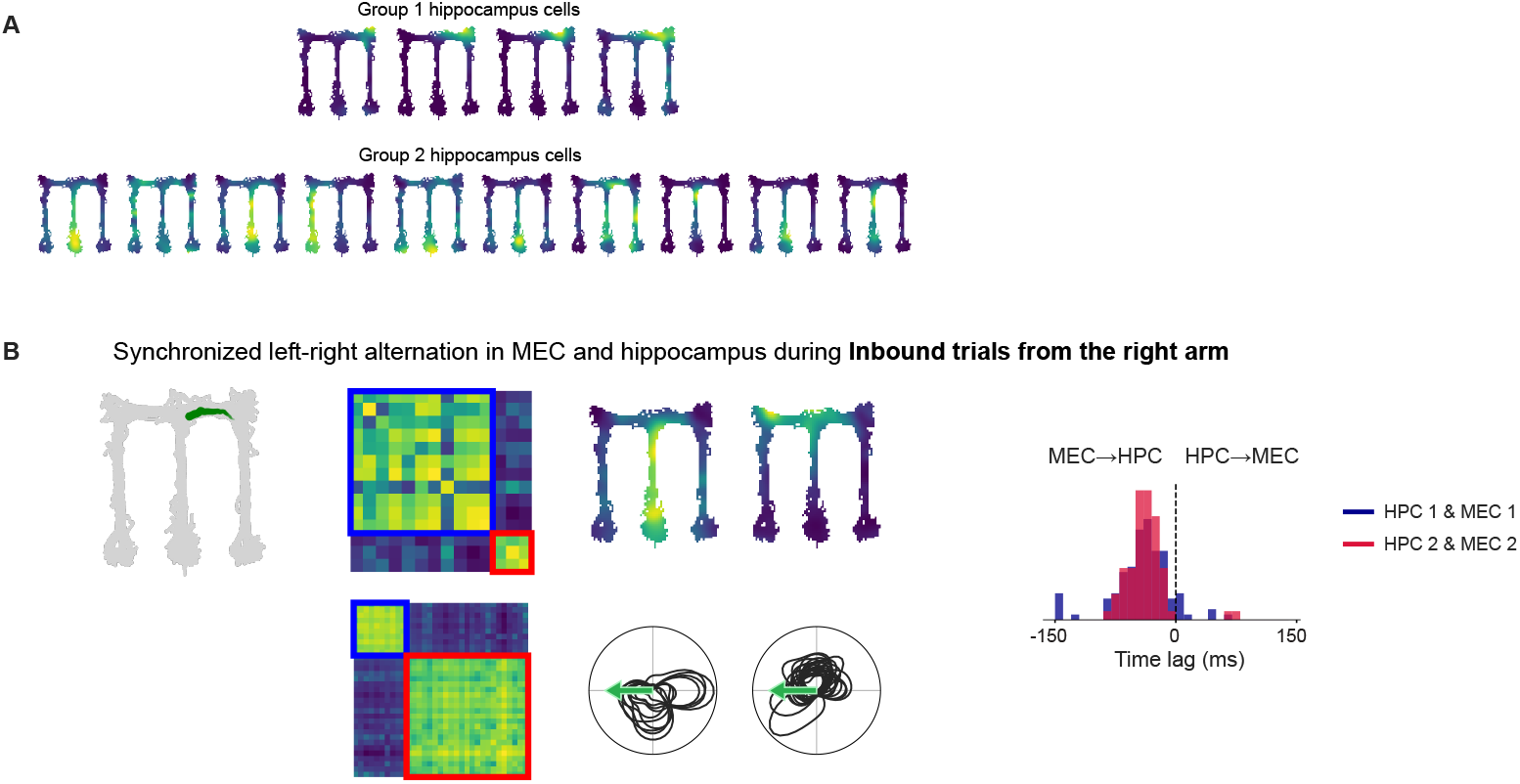
(**A**) Individual firing-rate maps of hippocampal cells belonging to the identified cluster. (**B**) Results for inbound trials when the animal was running from the right arm.

**Fig. S4.**
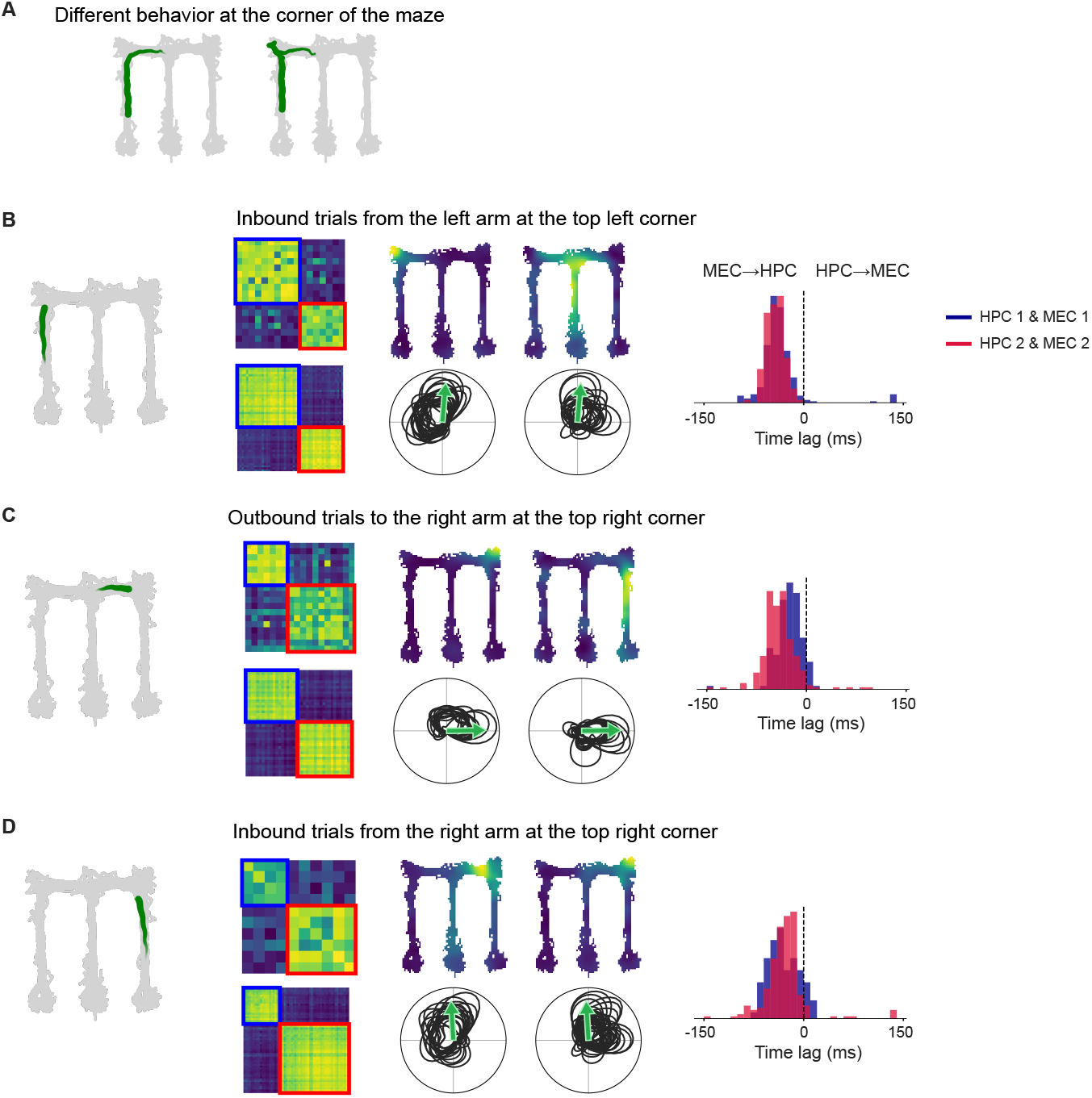
(**A**) Two example trajectories at the maze corner. The animal typically turns smoothly at the corner (left) but occasionally enters the dead end before turning (right). (**B**) Results for inbound trials from the left arm at the top-left corner. (**C**) Results for outbound trials to the right arm at the top-right corner. (**D**) Results for inbound trials from the right arm at the top-left corner

## Notes

### Competing Interest Statement

The authors have declared no competing interest.

### Summary of Updates

Update in the discussion section to reflect recent findings.

## References

1. W. B. Scoville, B. Milner, Loss of recent memory after bilateral hippocampal lesions. J. Neurol. Neurosurg. Psychiatry 20, 11–21 (1957).

2. D. S. Olton, J. T. Becker, G. E. Handelmann, Hippocampus, space, and memory. Behav. Brain Sci. 2, 313–322 (1979).

3. D. S. Olton, J. A. Walker, F. H. Gage, Hippocampal connections and spatial discrimination. Brain Res. 139, 295–308 (1978).

4. D. L. Schacter, D. R. Addis, D. Hassabis, V. C. Martin, R. N. Spreng, K. K. Szpunar, The future of memory: remembering, imagining, and the brain. Neuron 76, 677–694 (2012).

5. D. Hassabis, D. Kumaran, S. D. Vann, E. A. Maguire, Patients with hippocampal amnesia cannot imagine new experiences. Proc. Natl. Acad. Sci. U. S. A. 104, 1726–1731 (2007).

6. T. I. Brown, V. A. Carr, K. F. LaRocque, S. E. Favila, A. M. Gordon, B. Bowles, J. N. Bailenson, A. D. Wagner, Prospective representation of navigational goals in the human hippocampus. Science 352, 1323–1326 (2016).

7. D. R. Addis, A. T. Wong, D. L. Schacter, Remembering the past and imagining the future: common and distinct neural substrates during event construction and elaboration. Neuropsychologia 45, 1363–1377 (2007).

8. H. Petsche, C. Stumpf, G. Gogolak, The significance of the rabbit’s septum as a relay station between the midbrain and the hippocampus. I. The control of hippocampus arousal activity by the septum cells. Electroencephalogr. Clin. Neurophysiol. 14, 202–211 (1962).

9. B. Hangya, Z. Borhegyi, N. Szilágyi, T. F. Freund, V. Varga, GABAergic neurons of the medial septum lead the hippocampal network during theta activity. J. Neurosci. 29, 8094–8102 (2009).

10. M. Vandecasteele, V. Varga, A. Berényi, E. Papp, P. Barthó, L. Venance, T. F. Freund, G. Buzsáki, Optogenetic activation of septal cholinergic neurons suppresses sharp wave ripples and enhances theta oscillations in the hippocampus. Proc. Natl. Acad. Sci. U. S. A. 111, 13535–13540 (2014).

11. G. Dragoi, G. Buzsáki, Temporal encoding of place sequences by hippocampal cell assemblies. Neuron 50, 145–157 (2006).

12. J. O’Keefe, M. L. Recce, Phase relationship between hippocampal place units and the EEG theta rhythm. Hippocampus 3, 317–330 (1993).

13. N. Burgess, C. Barry, J. O’Keefe, An oscillatory interference model of grid cell firing. Hippocampus 17, 801–812 (2007).

14. W. E. Skaggs, B. L. McNaughton, M. A. Wilson, C. A. Barnes, Theta phase precession in hippocampal neuronal populations and the compression of temporal sequences. Hippocampus 6, 149–172 (1996).

15. D. J. Foster, M. A. Wilson, Hippocampal theta sequences. Hippocampus 17, 1093–1099 (2007).

16. A. S. Gupta, M. A. A. van der Meer, D. S. Touretzky, A. D. Redish, Segmentation of spatial experience by hippocampal θ sequences. Nat. Neurosci. 15, 1032–1039 (2012).

17. T. Feng, D. Silva, D. J. Foster, Dissociation between the experience-dependent development of hippocampal theta sequences and single-trial phase precession. J. Neurosci. 35, 4890–4902 (2015).

18. C. Liu, R. Todorova, W. Tang, A. Oliva, A. Fernandez-Ruiz, Associative and predictive hippocampal codes support memory-guided behaviors. Science 382, eadi8237 (2023).

19. K. D. Harris, J. Csicsvari, H. Hirase, G. Dragoi, G. Buzsáki, Organization of cell assemblies in the hippocampus. Nature 424, 552–556 (2003).

20. K. D. Harris, D. A. Henze, H. Hirase, X. Leinekugel, G. Dragoi, A. Czurkó, G. Buzsáki, Spike train dynamics predicts theta-related phase precession in hippocampal pyramidal cells. Nature 417, 738–741 (2002).

21. Y. Wang, S. Romani, B. Lustig, A. Leonardo, E. Pastalkova, Theta sequences are essential for internally generated hippocampal firing fields. Nat. Neurosci. 18, 282–288 (2015).

22. D. Robbe, G. Buzsáki, Alteration of theta timescale dynamics of hippocampal place cells by a cannabinoid is associated with memory impairment. J. Neurosci. 29, 12597–12605 (2009).

23. K. A. Bolding, J. Ferbinteanu, S. E. Fox, R. U. Muller, Place cell firing cannot support navigation without intact septal circuits. Hippocampus 30, 175–191 (2020).

24. M. J. Kahana, R. Sekuler, J. B. Caplan, M. Kirschen, J. R. Madsen, Human theta oscillations exhibit task dependence during virtual maze navigation. Nature 399, 781–784 (1999).

25. D. Arnolds, F. H. L. da Silva, J. W. Aitink, A. Kamp, P. Boeijinga, The spectral properties of hippocampal EEG related to behaviour in man. Electroencephalogr. Clin. Neurophysiol. 50, 324–328 (1980).

26. J. Winson, Loss of hippocampal theta rhythm results in spatial memory deficit in the rat. Science 201, 160–163 (1978).

27. G. Buzsáki, E. I. Moser, Memory, navigation and theta rhythm in the hippocampal-entorhinal system. Nat. Neurosci. 16, 130–138 (2013).

28. A. Johnson, A. D. Redish, Neural ensembles in CA3 transiently encode paths forward of the animal at a decision point. J. Neurosci. 27, 12176–12189 (2007).

29. A. M. Wikenheiser, A. D. Redish, Hippocampal theta sequences reflect current goals. Nat. Neurosci. 18, 289–294 (2015).

30. K. Kay, J. E. Chung, M. Sosa, J. S. Schor, M. P. Karlsson, M. C. Larkin, D. F. Liu, L. M. Frank, Constant Sub-second Cycling between Representations of Possible Futures in the Hippocampus. Cell 180, 552–567.e25 (2020).

31. C. Yu, Z. Ji, J. Ormond, J. O’Keefe, N. Burgess, Hippocampal theta sweeps indicate goal direction, bioRxiv (2025). 10.1101/2025.08.21.671551.

32. W. Tang, X. Mei, R. E. Harvey, E. Carbajal-Leon, T. Netzer, H. Chang, A. Oliva, A. Fernandez-Ruiz, Goal-directed hippocampal theta sweeps during memory-guided navigation, bioRxivorg (2025). 10.1101/2025.08.26.672489.

33. A. E. Comrie, E. J. Monroe, A. E. Kahn, E. L. Denovellis, A. Joshi, J. A. Guidera, T. A. Krausz, J. D. Berke, N. D. Daw, L. M. Frank, Hippocampal representations of alternative possibilities are flexibly generated to meet cognitive demands, bioRxivorg (2024). 10.1101/2024.09.23.613567.

34. W. Tang, J. D. Shin, S. P. Jadhav, Multiple time-scales of decision-making in the hippocampus and prefrontal cortex. Elife 10 (2021).

35. Z. Kurth-Nelson, T. Behrens, G. Wayne, K. Miller, L. Luettgau, R. Dolan, Y. Liu, P. Schwartenbeck, Replay and compositional computation. Neuron 111, 454–469 (2023).

36. M. G. Mattar, M. Lengyel, Planning in the brain. Neuron 110, 914–934 (2022).

37. M. E. Hasselmo, C. Bodelón, B. P. Wyble, A proposed function for hippocampal theta rhythm: separate phases of encoding and retrieval enhance reversal of prior learning. Neural Comput. 14, 793–817 (2002).

38. A. E. Comrie, L. M. Frank, K. Kay, Imagination as a fundamental function of the hippocampus. Philos. Trans. R. Soc. Lond. B Biol. Sci. 377, 20210336 (2022).

39. H. Davoudi, D. J. Foster, Acute silencing of hippocampal CA3 reveals a dominant role in place field responses. Nat. Neurosci. 22, 337–342 (2019).

40. M. I. Schlesiger, C. C. Cannova, B. L. Boublil, J. B. Hales, E. A. Mankin, M. P. Brandon, J. K. Leutgeb, C. Leibold, S. Leutgeb, The medial entorhinal cortex is necessary for temporal organization of hippocampal neuronal activity. Nat. Neurosci. 18, 1123–1132 (2015).

41. A. Fernández-Ruiz, A. Oliva, G. A. Nagy, A. P. Maurer, A. Berényi, G. Buzsáki, Entorhinal-CA3 dual-input control of spike timing in the hippocampus by theta-gamma coupling. Neuron 93, 1213–1226.e5 (2017).

42. B. Lasztóczi, T. Klausberger, Hippocampal place cells couple to three different gamma oscillations during place field traversal. Neuron 91, 34–40 (2016).

43. J. Suh, A. J. Rivest, T. Nakashiba, T. Tominaga, S. Tonegawa, Entorhinal cortex layer III input to the hippocampus is crucial for temporal association memory. Science 334, 1415–1420 (2011).

44. N. T. M. Robinson, J. B. Priestley, J. W. Rueckemann, A. D. Garcia, V. A. Smeglin, F. A. Marino, H. Eichenbaum, Medial entorhinal cortex selectively supports temporal coding by hippocampal neurons. Neuron 94, 677–688.e6 (2017).

45. T. Hafting, M. Fyhn, S. Molden, M.-B. Moser, E. I. Moser, Microstructure of a spatial map in the entorhinal cortex. Nature 436, 801–806 (2005).

46. R. J. Gardner, E. Hermansen, M. Pachitariu, Y. Burak, N. A. Baas, B. A. Dunn, M.-B. Moser, E. I. Moser, Toroidal topology of population activity in grid cells. Nature 602, 123–128 (2022).

47. T. Solstad, C. N. Boccara, E. Kropff, M.-B. Moser, E. I. Moser, Representation of geometric borders in the entorhinal cortex. Science 322, 1865–1868 (2008).

48. Ø. A. Høydal, E. R. Skytøen, S. O. Andersson, M.-B. Moser, E. I. Moser, Object-vector coding in the medial entorhinal cortex. Nature 568, 400–404 (2019).

49. S. S. Deshmukh, D. Yoganarasimha, H. Voicu, J. J. Knierim, Theta modulation in the medial and the lateral entorhinal cortices. J. Neurophysiol. 104, 994–1006 (2010).

50. A. Z. Vollan, R. J. Gardner, M.-B. Moser, E. I. Moser, Left-right-alternating theta sweeps in entorhinal-hippocampal maps of space. Nature 639, 995–1005 (2025).

51. A. Z. Vollan, R. Gardner, M.-B. Moser, E. Moser, Left-right-alternating theta sweeps in the entorhinal-hippocampal spatial map (v1), EBRAINS (2024); 10.25493/R5FR-EDG.

52. J. S. Taube, R. U. Muller, J. B. Ranck Jr, Head-direction cells recorded from the postsubiculum in freely moving rats. I. Description and quantitative analysis. J. Neurosci. 10, 420–435 (1990).

53. C. N. Boccara, F. Sargolini, V. H. Thoresen, T. Solstad, M. P. Witter, E. I. Moser, M.-B. Moser, Grid cells in pre- and parasubiculum. Nat. Neurosci. 13, 987–994 (2010).

54. D. Derdikman, J. R. Whitlock, A. Tsao, M. Fyhn, T. Hafting, M.-B. Moser, E. I. Moser, Fragmentation of grid cell maps in a multicompartment environment. Nat. Neurosci. 12, 1325–1332 (2009).

55. K. Mizuseki, A. Sirota, E. Pastalkova, G. Buzsáki, Theta oscillations provide temporal windows for local circuit computation in the entorhinal-hippocampal loop. Neuron 64, 267–280 (2009).

56. H. F. Ólafsdóttir, F. Carpenter, C. Barry, Coordinated grid and place cell replay during rest. Nat. Neurosci. 19, 792–794 (2016).

57. J. Huxter, N. Burgess, J. O’Keefe, Independent rate and temporal coding in hippocampal pyramidal cells. Nature 425, 828–832 (2003).

58. M. B. Zugaro, L. Monconduit, G. Buzsáki, Spike phase precession persists after transient intrahippocampal perturbation. Nat. Neurosci. 8, 67–71 (2005).

59. M. P. Brandon, A. R. Bogaard, N. W. Schultheiss, M. E. Hasselmo, Segregation of cortical head direction cell assemblies on alternating θ cycles. Nat. Neurosci. 16, 739–748 (2013).

60. S. P. Jadhav, C. Kemere, P. W. German, L. M. Frank, Awake hippocampal sharp-wave ripples support spatial memory. Science 336, 1454–1458 (2012).

61. A. Joshi, A. E. Comrie, S. Bray, A. Mankili, J. A. Guidera, R. Nevers, X. Sun, E. Monroe, V. Kharazia, R. Ly, D. A. Maya, D. Morales-Rodriguez, J. Yu, A. Kiseleva, V. Perez, L. M. Frank, Disruption of theta-timescale spiking impairs learning but spares hippocampal replay, bioRxivorg (2025) p. 2025.09. 15.675587.

62. J. Ormond, J. O’Keefe, Hippocampal place cells have goal-oriented vector fields during navigation. Nature 607, 741–746 (2022).

63. A. Z. Vollan, M. F. Schellenberger, M.-B. Moser, E. I. Moser, Attention-like regulation of theta sweeps in the brain’s spatial navigation circuit, bioRxiv (2026) p. 2026.01.27.702083.

64. Z. Ji, T. Chu, S. Wu, N. Burgess, A systems model of alternating theta sweeps via firing rate adaptation. Curr. Biol. 35, 709–722.e5 (2025).

65. J. Widloski, D. Theurel, D. J. Foster, Spontaneous alternation of place-cell sequences in the open field through spike frequency adaptation. Cell Rep. 44, 115475 (2025).

66. E. R. Wood, P. A. Dudchenko, R. J. Robitsek, H. Eichenbaum, Hippocampal neurons encode information about different types of memory episodes occurring in the same location. Neuron 27, 623–633 (2000).

67. L. M. Frank, E. N. Brown, M. Wilson, Trajectory encoding in the hippocampus and entorhinal cortex. Neuron 27, 169–178 (2000).

68. H. T. Ito, S.-J. Zhang, M. Witter, E. Moser, M. Moser, A prefrontal–thalamo–hippocampal circuit for goal-directed spatial navigation. Nature 522, 50–55 (2015).

69. C. R. Quirk, I. Zutshi, S. Srikanth, M. L. Fu, N. Devico Marciano, M. K. Wright, D. F. Parsey, S. Liu, R. E. Siretskiy, T. L. Huynh, J. K. Leutgeb, S. Leutgeb, Precisely timed theta oscillations are selectively required during the encoding phase of memory. Nat. Neurosci. 24, 1614–1627 (2021).

70. Y. Zhang, L. Cao, V. Varga, M. Jing, M. Karadas, Y. Li, G. Buzsáki, Cholinergic suppression of hippocampal sharp-wave ripples impairs working memory. Proc. Natl. Acad. Sci. U. S. A. 118, e2016432118 (2021).

71. T. Gedankien, J. Kriegel, E. Zabeh, D. McDonagh, B. Lega, J. Jacobs, Cholinergic blockade reveals a role for human hippocampal theta in memory encoding but not retrieval, bioRxivorg (2025). 10.1101/2025.05.12.653487.

72. T. M. George, M. Rastogi, W. de Cothi, C. Clopath, K. Stachenfeld, C. Barry, RatInABox, a toolkit for modelling locomotion and neuronal activity in continuous environments. Elife 13 (2024).

